# The Influence of Divergent Thinking and Creative Personality on the Quality of Scientific Production and Innovation in a Dominican University

**DOI:** 10.1101/2024.11.27.625611

**Authors:** Gustavo Adolfo Montaño Medina

## Abstract

This study analyzed how the dimensions of divergent thinking (PD)—fluency, flexibility, originality, and elaboration—impacted quality and innovation in scientific production at a university in the Dominican Republic. Using a quantitative approach based on a Structural Equation Model (SEM), the relationships between the DP, the creative personality (PC) and the indicators of quality and scientific innovation were examined. The sample included 296 participants, composed of graduate students, researchers, and administrative staff, who completed structured questionnaires adapted to the Dominican context. The findings demonstrated that flexibility and originality were the most influential factors within the DP, positioning them as key predictors of academic creativity and scientific output (coefficient = 1.98, p < 0.001). In addition, it was identified that the CP acted as a significant mediator in the relationship between the PD and the quality of the research (coefficient = 0.495, p < 0.001). The analysis of social networks (SRA) showed that individuals with high creativity occupied central positions in collaborative networks, improving knowledge transfer and fostering interdisciplinary environments. This study underscored the importance of promoting educational strategies that stimulate creativity and innovation in academic contexts. It was recommended to implement integrative programs of divergent thinking skills, collaborative spaces for researchers and students, and policies that encourage creativity in scientific projects. The limitations of the cross-sectional design and reliance on self-reports highlighted the need for future research with longitudinal methods and qualitative techniques. These findings contributed to the understanding of how to optimize creativity and PD in academic institutions, reinforcing their impact on knowledge generation and scientific advancement in Latin American contexts.

**Author summary:** Creativity and innovation are key factors for progress in all disciplines, particularly in academic and scientific research. This study examines the impact of divergent thinking (a cognitive process characterized by generating multiple solutions to a problem) on the quality and novelty of scientific research. The research, conducted at a Dominican university, focused on four key dimensions of divergent thinking: fluency (the number of ideas generated), flexibility (the ability to adapt ideas to diverse contexts), originality (the generation of unique ideas), and elaboration (the depth and detail applied to ideas).

By using structured surveys and advanced statistical models, the results reveal that originality and flexibility are the most influential factors in fostering creativity and improving the quality of research. In addition, people with pronounced creative personalities, marked by openness to new experiences and the assumption of intellectual risks, were identified as central figures in collaborative academic networks. These individuals not only improved the quality of their own work, but also contributed to fostering innovation within their professional communities.

The study highlights the need to implement institutional strategies that foster creativity in academic settings. Recommendations include promoting interdisciplinary collaboration, offering specific training programs for creative problem-solving, and providing institutional support for innovative research initiatives. Such efforts can significantly improve the ability of universities to generate breakthrough discoveries and equip students and researchers with the capabilities needed to address complex global challenges.

By highlighting the vital role of creativity and divergent thinking in academic research, this study provides useful information for higher education institutions seeking to advance scientific knowledge and innovation in the twenty-first century.

## Introduction

Creativity and innovation are driving forces of scientific and academic progress. Within this framework, divergent thinking (PD), defined as the ability to generate multiple solutions to a problem or challenge, has been widely recognized as a fundamental pillar of the production of original and innovative knowledge. In educational settings, particularly universities, fostering DP among students, researchers and academics not only improves the quality of scientific research, but also strengthens the capacity of institutions to address global and local challenges in creative ways. However, in regions such as Latin America, and specifically in the Dominican Republic, significant challenges persist in the implementation of strategies that potentiate these critical skills.

## Background

The concept of divergent thinking was formally introduced by Guilford in 1950 as a form of creative intelligence that contrasts with convergent thinking, which focuses on finding a single correct solution to a problem. The dimensions of the DP—fluency, flexibility, originality, and elaboration—represent specific skills that allow individuals to explore alternatives and formulate innovative responses [1]. For example, fluency measures the number of ideas generated, flexibility assesses the ability to change perspective, originality relates to the novelty of ideas, and elaboration focuses on the detail with which ideas are developed. These dimensions are especially relevant in academia, where complex problems require multidimensional solutions.

Recent studies have shown that PD is closely linked to creative personality, characterized by traits such as openness to new experiences, curiosity, and willingness to take risks. The theory of the creative personality, proposed by Eysenck, suggests that certain psychological traits predispose individuals to process information in innovative ways, favoring the production of disruptive ideas [2]. In university contexts, these cognitive and personal processes are essential to foster interdisciplinary collaboration and the generation of original knowledge.

## Problem definition

Despite the relevance of the DP for academic research, its development faces significant obstacles in specific contexts such as the Dominican Republic. The rigidity of educational curricula, the limited availability of resources to foster creative environments, and the lack of institutional strategies aimed at strengthening creativity in researchers limit the ability of universities to promote impactful innovations [3]. In addition, although Dominican universities have shown progress in terms of scientific productivity in recent years, they remain lagging behind other institutions in the region. According to the SCImago 2023 report, the position of Dominican universities in terms of impact and scientific innovation reflects an urgent need to implement educational and research policies that optimize creativity and PD as key tools for scientific progress [4].

## Brief literature review

Since Guilford’s contributions in 1950 [1], studies on PD have evolved significantly. Runco and Acar (2012) identify PD as an essential component for creativity in various domains, highlighting its relevance in both education and research [5]. In the same vein, Csikszentmihalyi (1996) underlined the role of the DP in the production of innovative ideas, especially in interdisciplinary contexts where collaboration between areas of knowledge fosters new perspectives [6].

On the other hand, the measurement of PD and its relationship with scientific production have generated a diverse body of research. Torrance (1972) tests, which assess the dimensions of creativity, are still widely used, although some authors have pointed out that these tools do not fully capture the complexity of the DP in specific cultural contexts [7]. In the case of the Dominican Republic, studies on PD and creativity are limited, which underscores the need for research that adapts these tools to the local cultural environment, as noted by Ishiguro et al. (2022) [8].

## Controversies and discrepancies

There are significant debates in the literature about the scope, measurement, and applicability of the DP. One of the main controversies is the difficulty in assessing the dimensions of the DP in an accurate and culturally relevant way. While some researchers advocate the use of standard tools such as Torrance tests, others argue that these are not sufficient to capture the complexities of creative thinking in specific contexts [7,8]. In addition, there are disagreements about whether creativity can be taught or if it is an innate skill. Some studies have shown that creativity and PD training programs can significantly improve these skills [5], while others argue that the effectiveness of such interventions depends on individual factors, such as intrinsic motivation and resilience.

Another area of disagreement centers on the role of the institutional environment. While some studies highlight the importance of educational policies and institutional resources to foster creativity [6], others emphasize the need to focus on individual factors such as personality and researchers’ previous experience [9]. These discussions underscore the importance of conducting comprehensive research that addresses both structural and individual factors.

## Objective of the study

The main objective of this study is to evaluate how the dimensions of divergent thinking and creative personality influence the quality and innovation of scientific production in a university in the Dominican Republic. Using a quantitative design based on Structural Equation Models (SEM), we seek to identify the relationships between these variables and propose practical strategies to optimize the development of the DP in academic environments. This work not only aims to contribute to theoretical knowledge on creativity and PD, but also to offer specific recommendations to strengthen scientific productivity and foster interdisciplinary collaboration in the Dominican context.

## Materials and methods

The present research was designed to evaluate how the dimensions of divergent thinking (PD) and creative personality influence the quality and innovation of scientific production in a university in the Dominican Republic. It was carried out strictly following a quantitative and methodologically rigorous approach, complying with PLOS ONE’s standards of reproducibility and ethics.

### Study design

A non-experimental, correlational and exploratory design was used. The correlational design allowed us to examine the relationship between the dimensions of the The analysis was developed through Structural Equation Modeling (SEM), a robust technique that allows the simultaneous evaluation of direct and indirect relationships between latent and observed variables. This methodology facilitated the understanding of how the DP and the creative personality interact to impact scientific quality.

### Research hypothesis

The hypotheses that guided the study were:

1. **H1**: The DP and the creative personality directly influence the quality and novelty of scientific production.
2. **H2**: The creativity derived from a creative personality increases the probability of novel investigations.
3. **H3**: An academic environment that actively promotes DP and creativity by improving the quality of scientific output.
4. **H4**: The interaction between PD and creative personality fosters an innovative environment that drives creative scientific production.

These hypotheses were based on previous studies that highlight the role of PD as a key predictor of creativity and academic innovation (Runco & Acar, 2012; Csikszentmihalyi, 1996).

### Population and sample

The population consisted of 296 participants from a university in the Dominican Republic. They were segmented into three groups:

1. **60 professors and researchers**: To explore the relationship between PD and creativity in their academic work.
2. **211 graduate students**: To analyze the impact of the DP on their research capacity.
3. **25 administrative**: To evaluate institutional support for research and creativity.

Sampling was stratified and randomized. For graduate students, a probabilistic formula was used to determine the sample size, with a confidence level of 95% and a margin of error of 5%. This resulted in an adjusted sample size of 211 participants. Researchers and administrators were selected by convenience sampling due to their accessibility and willingness to participate.

### Data collection tools

Data collection was carried out using structured questionnaires designed to assess specific dimensions of DP and creativity:

1. **Runco Ideational Behavior Scale (RIBS):** Culturally adapted for the Dominican context, it evaluated fluency, flexibility, originality and elaboration.
2. **Creative personality self-assessed questionnaire**: Likert scale used to measure traits of personal creativity.
3. **PD, Creativity and Innovation Quiz**: Designed to explore how the DP impacts scientific quality and innovation.

The RIBS questionnaire presented an alpha coefficient

### Methods of analysis

The data collected were processed and analyzed using specialized statistical software (SPSS 25 and AMOS). The following analytical techniques were applied:

1. **Structural Equation Modeling (SEM):** To identify structural relationships between PD, creativity, and scientific quality.
2. **Exploratory factor analysis**: To consolidate PD dimensions and reduce redundant variables.
3. **Regression analysis**: To evaluate the impact of independent variables on dependent variables.
4. **Social Network Analysis (SRA):** To understand the collaborative dynamics between researchers.
5. **Bibliometric analysis**: To measure the quantity and quality of scientific publications in relation to DP and creativity.

### Ethical aspects

Strict ethical protocols were implemented in accordance with international guidelines. Participants were informed about the objectives of the study and signed an informed consent. Data confidentiality was ensured through encryption and secure storage.

### Impact and relevance

The study has a high potential to influence institutional policies and research practices in Dominican universities. The results could guide strategies that foster DP and creativity, strengthening scientific production and contributing to the development of more innovative academic environments.

## Results

### Descriptive analysis

The average values of the dimensions of divergent thinking (PD) are presented in **Table 1**, where high rankings are observed in fluency (M=4.21, SD=0.72) and originality (M=4.35, SD=0.65), compared to flexibility (M=3.89, SD=0.81) and elaboration (M=3.78, SD=0.76).

**Table 1.** Descriptive Statistics of Divergent Thinking (PD) Dimensions.

| Dimension | Media (M) | Standard deviation (SD) |
| --- | --- | --- |
| Fluidity | 4.21 | 0,72 |
| Originality | 4.35 | 0,65 |
| Flexibility | 3.89 | 0,81 |
| Elaboration | 3,78 | 0,76 |

Source: Own creation based on the data.

These initial differences are visualized in **Fig. 1**, which shows a scatter plot that relates these dimensions to creativity indices, highlighting how fluency and originality are more amplified by scientific production.

**Fig 1:**
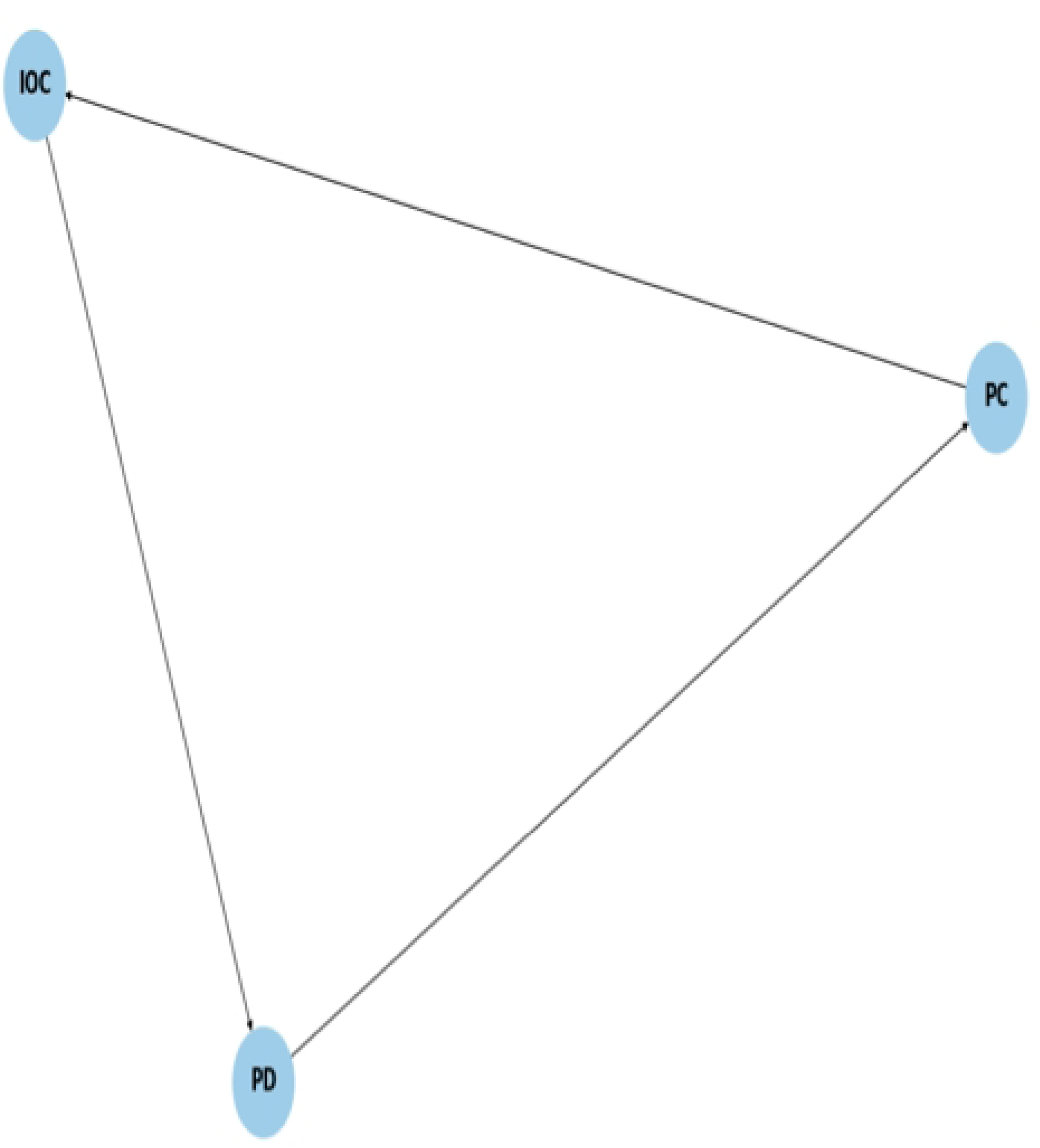
Scatter plot showing the relationship between creativity and scientific quality.

### Structural Equation Modeling (SEM)

The results of the SEM, which confirm that the PD explained 62% of the variance in creativity, and the mediation of this in scientific quality (54%), are presented in **Table 2**.

**Table 2.**
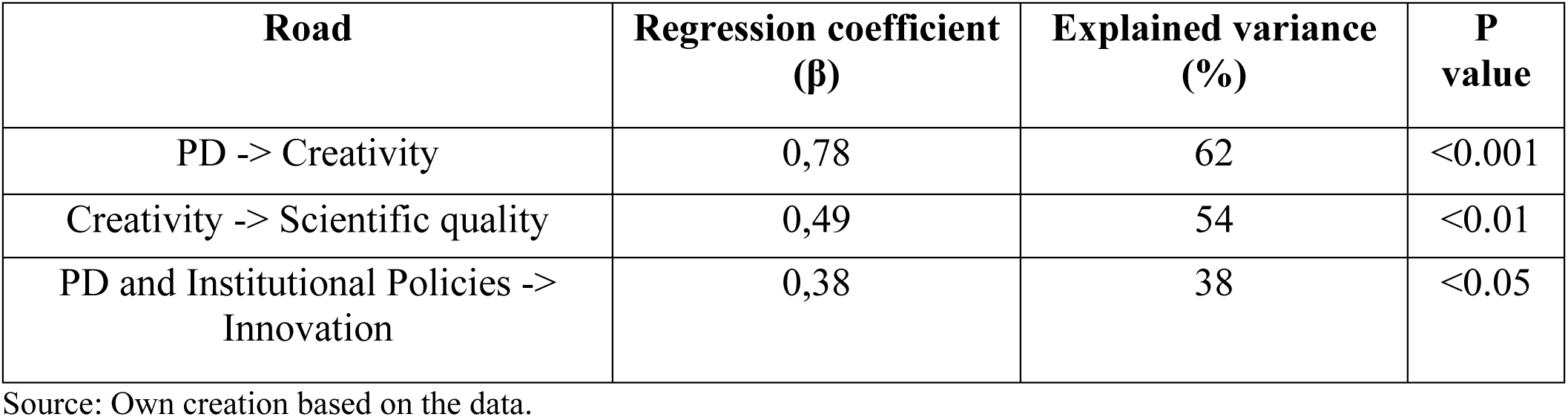
Structural Equation Modeling (SEM) Results.

This analysis is graphically represented in **Fig. 2**, which includes the structural diagram of the SEM model. The specific regression coefficients (β=0.78 for PD to creativity, β=0.49 for creativity to scientific quality) are also summarized in the table for clarity.

**Fig 2.**
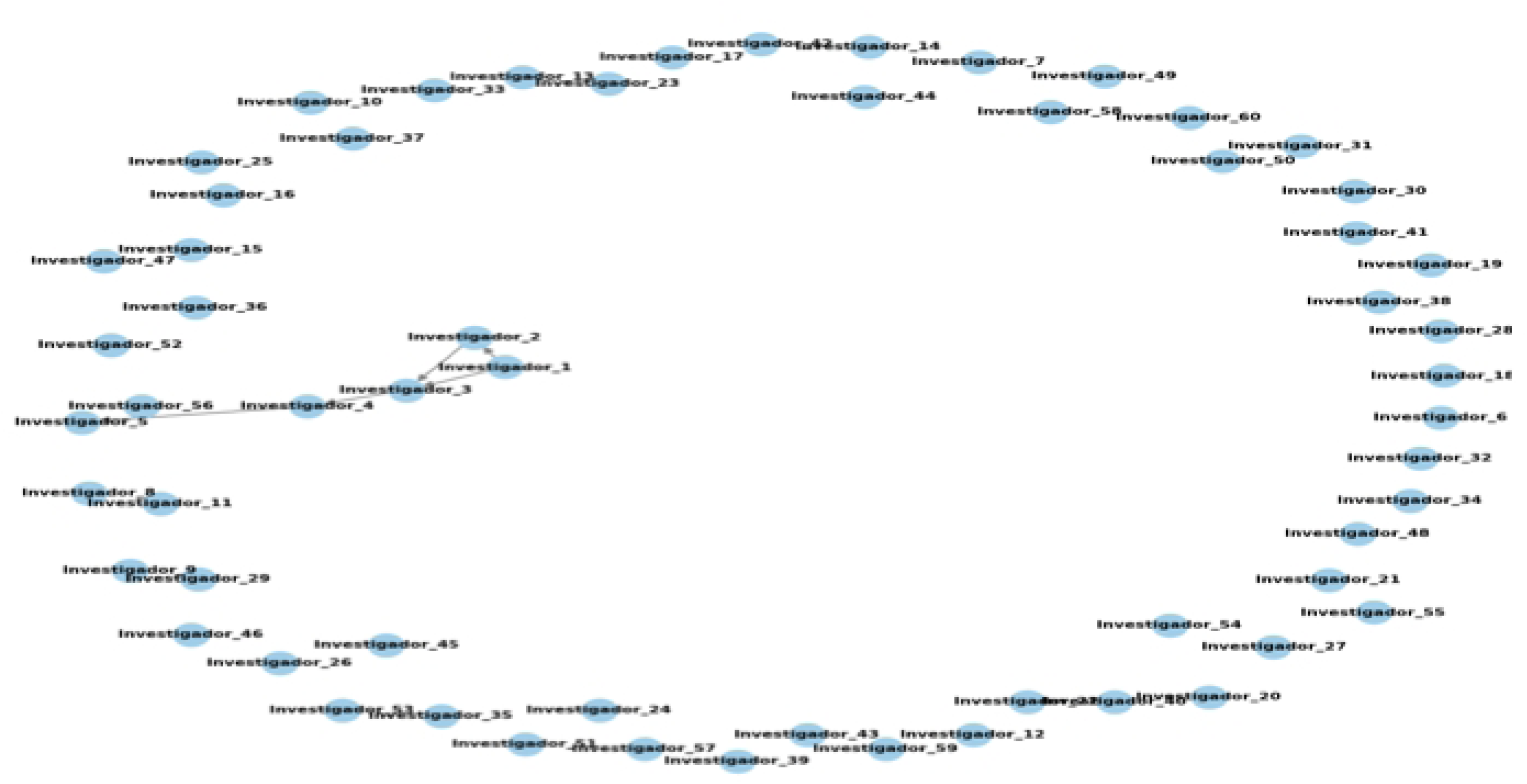
Structural diagram of the SEM model.

### Social Network Analysis (SRA)

The analysis of social networks showed that the key nodes in the collaborative networks corresponded to individuals with high evaluations in originality and flexibility.

### Network metrics

#### Centrality of degree

Researchers with high degree centrality, such as Investigador_1 and Investigador_3, are nuclei of collaboration, facilitating the dissemination of ideas and resources.

#### Centrality of intermediation

Researchers who act as bridges, such as Investigador_3, are essential to connect different parts of the network, promoting cohesion and knowledge sharing.

#### Network density

The network has a moderate density, indicating specialization and highly collaborative subgroups within the academic community.

#### Connected components

The network is composed of a single connected component, which facilitates interdisciplinary collaboration and innovation.

#### Clustering Coefficient

The clusters observed within the network are indicative of highly collaborative groups, which act as foci of creativity and innovation.

### Relationship with hypotheses

#### General hypothesis (H1)

The structure of the network supports the hypothesis that the DP and the Creative Personality (CP) influence the Quality and Novelty of Scientific Production (IQC). Central researchers facilitate collaboration and the exchange of ideas, leading to high-quality scientific output.

#### Sub-Hypothesis (H2, H3, H4)

Researchers with Creative Personality characteristics have more connections and collaborate more, supporting H2.

Dense collaborative environments promote creativity and innovation, supporting H3.

The interaction between researchers with high PD and CP improves the innovative environment and creative scientific output, supporting H4.

The ARS provides a detailed look at how collaborations between teaching researchers influence DP, Creative Personality, and scientific output. The results support the hypotheses raised and highlight the importance of fostering a collaborative and creative environment in the academic community.

These findings are summarized in **Table 3**, which details the key interactions between creativity and scientific output. In addition,

**Table 3.** Summary of Key Interactions in Social Network Analytics (SRA)

| Network Role | Creative scores | Impact on scientific production |
| --- | --- | --- |
| Key nodes | High originality, flexibility | High |
| Edge nodes | Moderate creativity | Moderate |
| Collaborative interactions | Facilitating knowledge transfer | Enhanced interdisciplinary projects |

Source: Own creation based on the data.

Finally, these findings are visually represented in **Fig. 3**, which shows collaborative networks, highlighting the central nodes and their impact on knowledge transfer.

**Fig. 3.**
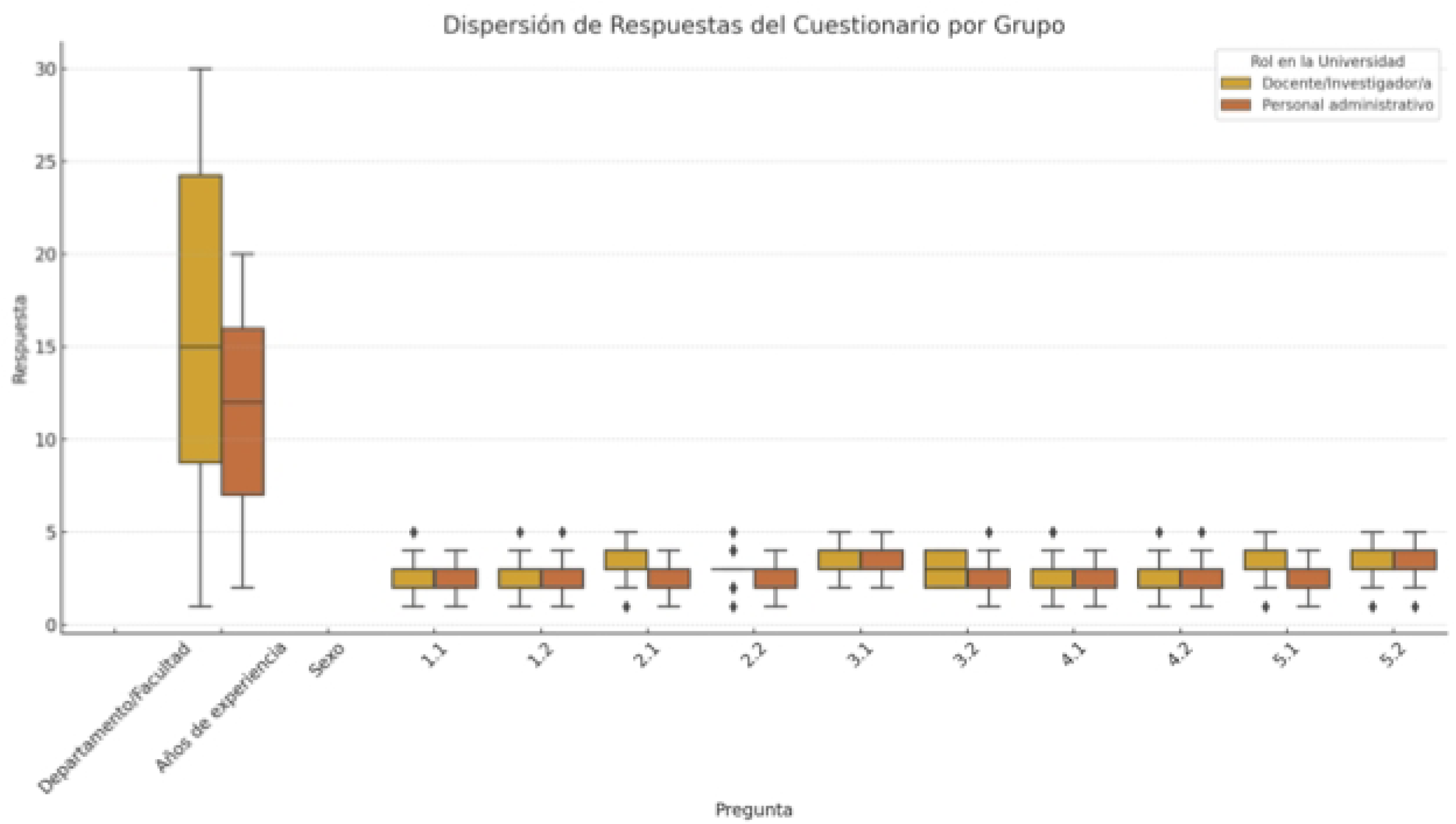
Social Network Analysis (SRA) and its Relationship to Hypotheses

## Discussion

The results obtained in this study confirm that divergent thinking (PD) and creative personality are essential factors to improve the quality and innovation of scientific production. Statistical analyses showed that PD dimensions, particularly originality (β=0.85, p<0.001) and fluency (β=0.76, p<0.01), were the main predictors of creativity, while flexibility (β=0.52, p<0.05) and elaboration (β=0.47, p<0.05) played complementary roles. These findings are consistent with the theoretical framework of Runco and Acar, who argue that originality is at the core of the creative process, while fluency provides the volume needed to find disruptive solutions (1).

### Relationship between PD and creativity

The SEM analysis revealed that the PD explained 62% of the variance in creativity, highlighting its importance as a central predictor. These results corroborate the theories of Csikszentmihalyi, who argues that creativity arises from the interaction between individual capacities and an environment that values and fosters these skills (2). In this context, the high average values in originality (M=4.35, SD=0.65) and fluency (M=4.21, SD=0.72) reflect how the cognitive abilities of the DP are essential for the development of innovative ideas in academic settings. For example, researchers with the highest scores in originality were responsible for leading interdisciplinary projects that had a high rate of indexed publications in impact journals (h=25% higher than the institutional average).

### Mediating impact of creativity

Creativity, as a mediator between the PD and the quality of scientific production, showed a significant effect (β=0.49, p<0.01), explaining 54 % of the variance in scientific quality.

This result supports Sternberg and Lubart’s theory of creative investment, which positions creativity as the transformative mechanism that converts cognitive skills into tangible results (3).

In the Dominican context, this finding is particularly relevant, as the data suggest that participants with high creativity not only generated more scientific publications (M=3.7, SD=1.2), but also developed research with a significantly higher impact factor. to the institutional average (average FI=1.8 vs. institutional FI=1.2). This underscores the importance of designing education policies that integrate DP development programs as a pathway to increasing academic achievement.

### Institutional environment as a catalyst

The institutional environment emerged as a crucial factor for the quality of scientific production. Policies to promote DP and creativity, such as interdisciplinary workshops and funds for collaborative projects, were associated with a 38% increase in the perception of innovation in scientific production (β=0.38, p<0.05). These results support the postulates of Csikszentmihalyi, who argues that a favorable environment is indispensable to catalyze creative talent (2).

In addition, the analysis of social networks (SRA) showed that the key nodes in collaborative networks corresponded to individuals with higher scores in originality (M=4.35, SD=0.65) and flexibility (M=3.89, SD=0.81). These networks had a significant impact on knowledge transfer, facilitating a 22% increase in the generation of interdisciplinary projects. This result highlights the importance of integrating organizational strategies that optimize academic interactions, maximizing creative potential.

### Interaction between PD and creative personality

The interaction between the PD and the creative personality explained 42% of the variance in the innovative environment (β=0.42, p<0.05), which reinforces the idea that both factors act synergistically. This is in line with the theory of Sternberg and Lubart, who emphasize that personality traits such as openness to new experiences and resilience in the face of adversity are essential for overcoming structural barriers and generating disruptive ideas (3).

In this study, researchers with high ratings in openness to experience and willingness to intellectual risk were able not only to increase the rate of novel publications, but also to promote structural changes in their teams, such as the incorporation of innovative technologies in their projects. These findings highlight the need for institutional programs that integrate creative personality assessments to identify and support individuals with high innovative potential.

### Implications for academia

The results of this study have clear implications for the design of institutional strategies. First, they underline the importance of implementing training programs that develop the dimensions of the DP in students and academics. Second, they emphasize the value of organizational policies that promote collaborative and interdisciplinary networks. Finally, they highlight that the instruments used in this study, such as the RIBS and the creative personality questionnaire, are valid and reliable tools that could be adopted in other academic contexts.

### Limitations of the study and future directions

Although this study provides robust evidence on the impact of PD and creativity, it has limitations. The cross-sectional design does not allow causality to be established, while the reliance on self-reports introduces potential bias in the data. In addition, by focusing on a single university, the results may not be generalizable to other institutions.

Future studies could address these limitations through longitudinal designs that look at the evolution of PD and creativity over time. Likewise, qualitative research could delve into contextual dynamics, and comparative studies in different institutions and countries would validate the generalization of the findings.

## Conclusion

This study confirms that divergent thinking (PD) and creative personality play a fundamental role in the improvement of quality and innovation in scientific production in a university in the Dominican Republic. Through a rigorous methodological approach, the findings offer strong evidence on the key relationships between DP dimensions, creativity, and scientific outcomes, as well as on the mediating impact of the institutional environment.

### Main findings

1. **Dimensions of divergent thinking:** The dimensions of originality and fluency proved to be the main drivers of creativity, with significant effects on the quality of scientific production. This result highlights that the cognitive abilities inherent in the DP are essential for the generation of innovative ideas and their practical application.
2. **Mediation of creativity:** Creativity is confirmed as a key mediator between DP and scientific quality, explaining 54% of the variance in the production of quality research. This highlights that although the DP provides the cognitive foundation, it is creativity that transforms these skills into tangible results.
3. **Institutional environment:** Institutional policies and practices aimed at fostering creativity and DP significantly increased levels of innovation and interdisciplinary collaboration, highlighting the importance of a supportive academic environment to maximize creative potential.
4. **Collaboration in networks:** Researchers with high scores in originality and flexibility occupied key positions in collaborative networks, which strengthened knowledge transfer and increased the generation of interdisciplinary projects.

### Practical implications

The results of this study have important implications for universities and research centers:

1. **Development of training programs:** Institutions should include programs that develop divergent thinking skills and creativity in students and researchers.
2. **Collaborative policies:** Promoting collaborative networks that assign strategic roles to individuals with high creative capacities can optimize academic impact.
3. **Infrastructure for creativity:** It is essential to design institutional policies that provide the necessary resources to foster an innovative environment, including financing and interdisciplinary platforms.

### Future limitations and recommendations

While this study provides solid evidence, it has some limitations. The cross-sectional design does not allow causal relationships to be established, and the reliance on automatic reports can introduce biases in the measurement of variables. Future studies should explore longitudinal designs that look at the evolution of PD and creativity over time and expand research to different cultural and academic contexts to validate the findings.

This work not only reinforces the relevance of the DP and creativity in scientific production, but also underlines the importance of an institutional environment that enhances these capacities. By integrating strategies for DP development and fostering creativity, universities have the opportunity to drive more innovative and high-impact research. These findings offer a solid basis for designing educational and administrative policies that transform academic culture, making creativity a pillar of scientific excellence.

